# Single and Multi-trait Genomic Prediction for agronomic traits in *Euterpe edulis*

**DOI:** 10.1101/2022.09.19.508517

**Authors:** Guilherme Bravim Canal, Cynthia Aparecida Valiati Barreto, Francine Alves Nogueira de Almeida, Iasmine Ramos Zaidan, Diego Pereira do Couto, Camila Ferreira Azevedo, Moysés Nascimento, Marcia Flores da Silva Ferreira, Adésio Ferreira

## Abstract

Popularly known as juçaizeiro, *Euterpe edulis* has been gaining prominence in the fruit growing sector and has demanded the development of superior genetic materials. As it is a native species and still little studied, the application of more sophisticated techniques can result in higher gains with less time. Until now, there are no studies that apply genomic prediction for this crop, especially in multi-trait analysis. In this sense, this study aimed to apply new methods and breeding techniques for the juçaizeiro, to optimize this breeding program through the application of genomic selection. This data consisted of 275 juçaizeiro genotypes from a population of Rio Novo do Sul-ES, Brazil. The genomic prediction was performed using the multi-trait (G-BLUP MT) and single-trait (G-BLUP ST) models and the selection of superior matrices was based on the selection index of Mulamba and Mock. Similar results for predictive ability were observed for both models. However, the G-BLUP ST model provided greater selection gains when compared to the G-BLUP MT. For this reason, the genomic estimated breeding values (GEBVs) from the G-BLUP ST were used to select the six superior genotypes (UFES.A.RN.390, UFES.A.RN.386, UFES.A.RN.080, UFES.A.RN.383, UFES.S.RN.098, and UFES.S.RN.093), to provide superior genetic materials for the development of seedlings and implantation of productive orchards, which will meet the demands of the productive, industrial and consumer market.

**Key message:** In the first genomic selection study for *Euterpe edulis*, substantial gains for multiple traits of fruit production was reported. This is a key factor for the sustainable use of the species in the Atlantic Forest.

## 1. Introduction

Native to the Atlantic Forest, *Euterpe edulis*, popularly known as juçaizeiro, has shown great economic potential (de Oliveira Maciel et al., 2019; Schulthess et al., 2016; Silva et al., 2018). Its fruits are used for the industrial production of processed pulp, which is similar to açaí (Silva et al., 2018). Given its productive potential and wild *status*, there is a demand for superior genetic materials to use the species as a new crop. However, due to a series of intrinsic characteristics, such as, the wild state of the species, slow development, difficulty in the practice of controlled pollination and propagation exclusively via seminal, classical breeding practices may not be sufficient to lead to satisfactory genetics gains in the short time currently required.

Within breeding programs, the genome-wide selection (GWS) proposed by Meuwissen et al. (2001), seeks through its techniques, such as G-BLUP (Genomic Best Linear Unbiased Prediction), the use of knowledge of genomic relationship to increase selective accuracy (Alkimim et al., 2020; de Resende et al., 2015). The application of the GWS in a natural population open-pollinated, as the base population of improvement of the juçaizeiro species, in which the degree of relationship between the individuals is unknown, is scarce in the literature. However, several studies were conducted on forest species from open-pollinated populations with knowledge of families (Bush and Thumma, 2013; El-Dien et al., 2018; El-Kassaby et al., 2011; Gamal El-Dien et al., 2016; Klápště et al., 2018) and support the assumption of increased accuracy with GWS application in a natural population without knowledge of relationship structure. This is due to the fact that, in these works, the use of the genomic relationship matrix (G) corrects the unrealistic priori that, in a given family, all individuals share the same genetic similarity with each other. In addition to the above, the accuracy of genomic prediction is generally higher, especially for breeding populations with superficial genealogy and disconnected families (Thavamanikumar et al., 2020).

Genomic prediction based on single traits has become a widely applied procedure among breeders after reducing genotyping costs (Jarquín et al., 2019; Massman et al., 2013; Sousa et al., 2019), mainly using the G-BLUP method. However, this model may not satisfactorily reflect the interactive complexities between the analyzed traits, because it does not capitalize on the flows of information between traits through available information on genetic covariances (Gaire et al., 2022). This covariances are generated through shared genetic influence (pleiotropy) and/or by the non-random association between the alleles (linkage disequilibrium), which, consequently, are responsible for generating complex relationships between quantitative characters (Lynch and Walsh, 1998). Therefore, the multi-trait approach (G-BLUP MT) applied to GWS has been highlighted for being able to combine information and capture the effect of association between traits in order to predict genetic values more accurately (Bhatta et al., 2020; Gaire et al., 2022; Lado et al., 2018; Sapkota et al., 2020).

Currently, for the juçaizeiro, the techniques and approaches applied to its scarce existing breeding programs are simple. In general, the selection of the best genotypes are carried out in natural populations, based on data from a single year and with a mass phenotypic selection technique, due to the absence of experimental fields with adult representatives of the species, without pre-established delineation and lack of knowledge of relationship between the individuals. In this sense, the selective bias can compromise the success of the species as a crop, by indicating low productivity genetic materials. Therefore, the present work aim to apply new methods and breeding techniques for juçaizeiro, to optimize the breeding programs through the application of genomic selection. This study was carried out in a natural open-pollinated population containing 275 genotypes, with no knowledge of relationship between individuals in order to select the genotypes with the greatest potential to serve the productive and industrial sector; and to compare the efficiency between multi-trait (G-BLUP MT) and single-trait (G-BLUP ST) models.

## 2. Materials and Methods

### 2.1. Field experimental conduction

Experimental evaluations were carried out from 2018 to 2021 in the municipality of Rio Novo do Sul in the state of Espírito Santo, Brazil (Fig. 1). The 275 genotypes evaluated were selected for good phytosanitary and physiological conditions and full reproductive age. The genotypes are located in a commercial plantation belonging to two companies that process juçaizeiro fruit pulp, Açaí juçara® and Bonalotti®. The plantation was formed by a mixture of individuals that emerged by spontaneous development, through the action of dispersers, and the enrichment by the owners by the sowing of seeds in the area. Commercial planting does not have pre-defined spacing and basically no productive management treatment, only mowing at harvest time. Consequently, the field evaluations followed a blank design, with initial unknown relationship by descent between the genotypes.

**Fig. 1.**
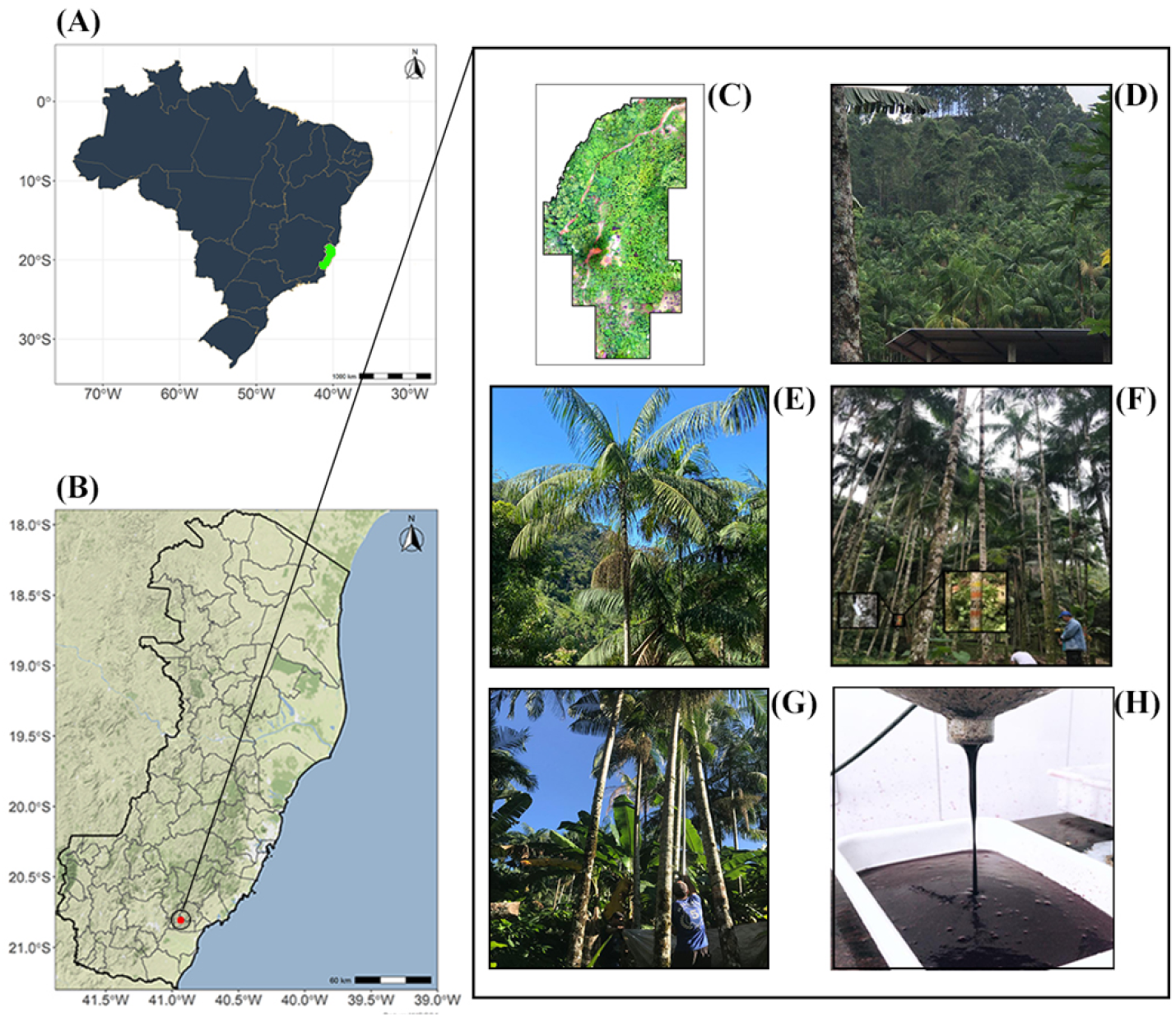
Geographical location of the experimental field where the *Euterpe edulis* matrices are located. Map generated with R free environment *software*. (A) Delimitation of Brazil, in green, the geographic location of the state of Espírito Santo; (B) Graphic representation of the state of Espírito Santo, in red, the location of the municipality of Rio Novo do Sul; (C) Orthomosaic of the experimental area; (D) Image referring to a fragment of the experimental planting; (E) Juçaizeiro (*Euterpe edulis*) from experimental planting in productive phase; (F) Registration of genotypes identification in field; (G) Juçaizeiro (*Euterpe edulis*) fruits harvest; (H) Processed pulp of juçaizeiro (*Euterpe edulis*) fruits.

### 2.2. Phenotyping of plants

In the years 2018 to 2021, the number of bunches per plant (NB) was evaluated by visually counting the number of bunches with fruits of each matrix plant. In the years 2018, 2019 and 2021, the mass of fruits per bunch (MFB) (kg), rachis length (RL) (cm), equatorial diameter of fruits (EDF) (mm) and pulp yield (PY) (%) were evaluated. For the evaluation of MFB, in all evaluations, the same trained professional performed the determination of the harvest point of the bunches. The harvest of each matrix was carried out when the fruit maturation point reached the stage used for fruit processing in the industry. Each year, one bunch per plant was harvested, the fruits were separated from the rachillas, and they were weighed on a scale with an accuracy of 0.1 g. For RL, after separating the fruits, the length of the rachis of the inflorescence was measured with a tape measure.

A sample of fruits was taken from each matrix, packed in properly identified plastic bags and transported to the Plant Biometry laboratory at the Federal University of Espírito Santo, where morphometric evaluations of fruits and seeds were carried out in a completely randomized design. For EDF, data were measured in millimeters (mm) obtained using a 6” digital caliper (Zaasprecision®), performed on five fruits individually, as recommended by Marçal et al. (2015) who observed high repeatability values for these characteristics. The authors concluded that five measurements are necessary to carry out the measurement to reach coefficients of determination of 95%, measurements above this amount increase costs and evaluative time, bringing little additional information to the works.

The PY was estimated by the following relationship:

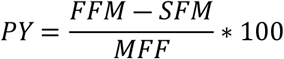

where FFM is fruit fresh mass and SFM is seed fresh mass, measured by weighing four replicates of 25 fruits and seeds, using an analytical balance (0.0001g).

### 2.3. Genomic DNA extraction

Genomic DNA was obtained from leaf samples of the genotypes under study. Extraction was performed using the cetyltrimethylammonium bromide or CTAB method by Doyle (1990) with modifications (Carvalho et al., 2020).

DNA concentrations and integrity were estimated using a NanodropTM 2000 spectrophotometer (Thermo Scientific). DNA quality was verified on 0.8% agarose gel. DNA genotypes prepared for genotyping using the DArTseqTM methodology were sent to the Service of Genetic Analysis for Agriculture (SAGA) in Mexico for high-throughput genotyping using the DArTseqTM technology

The genome representation of the 275 genotypes was obtained from the reduction of DNA complexity using two restriction enzymes, HpaII and Msel, and the ends of the cleaved fragments were linked to a code adapter and a common adapter to identify each sample. The fragments were amplified by PCR reaction; subsequently, equimolar amounts of amplification products from each sample of the 96-well microtiter plate were pooled, purified and quantified, then sequenced on the Illumina Novaseq 6000 System platform. The sequences were analyzed using Dartsoft14, an automated genomics data analysis program and DArTdb, a laboratory management system, developed and patented by DArT Pvt. Ltd. (Australia), generating SNP marker data.

### 2.4. Quality control of molecular markers

The dataset of codominant markers of the SNP type was submitted to quality control analysis in the R (R Core Team, 2021). The quality parameters used were *Call Rate* (CR) of 90% and *Minor Allele Frequency* (MAF) of 1%. After quality control, the marker dataset reduced by 81.75% from 44,457 markers to 8,112.

### 2.5. Phenotypic data analysis

The phenotypic values were corrected for the effect of years in the R software according to the correction proposed by (Fanelli Carvalho et al., 2020). The linear model used was:

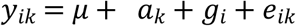

where *y_ik_* is the phenotypic value for the genotype *i* and the year *k*; *μ* is the population mean; *a_k_* is the fixed effect of the *k*^th^ year; *g_i_* is the fixed effect of the *i*^th^ genotype; and *e_ik_* is the random effect of residual, with 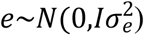. The empirical best linear unbiased estimates (eBLUEs) was calculated for each traits individually and their values were obtained by 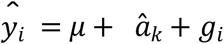.

### 2.6. Genomic prediction (GP)

Genomic prediction was performed using the multi-trait (G-BLUP MT) and single-trait (G-BLUP ST) models. For this, was used the SOMMER package version 3.4 [26], implemented in the R software (R Core Team, 2021). The G-BLUP MT model for the prediction of the genomic estimated breeding values (GEBV) of the individuals used was:

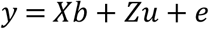

Where, y is the vector of previously estimated eBLUES and structured as *y* = [*y*_1_ *y*_2_ … *y_n_*]’; *y*_1_, *y*_2_ …, *y_n_* is the vector of observations for each characteristic; *b* is the vector of means of each characteristic structured as *b* = [*b*_1_ *b*_2_ … *b_n_*]’ and with X incidence matrix; *u* is vector of individual additive genomic genetic values of each trait structured as *u* = [*u*_1_ *u*_2_ … *u_n_*]’ with Z incidence matrix, with variance structure given by 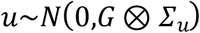, where *G* is the genomic relationship matrix between individuals for additive effects, *∑_u_* is the additive genetic covariance matrix and ⊗ denotes the Kronecker product; *e* is the random error vector with 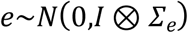 where *∑_e_* is the residual covariance matrix.

Covariance matrices can be written as:

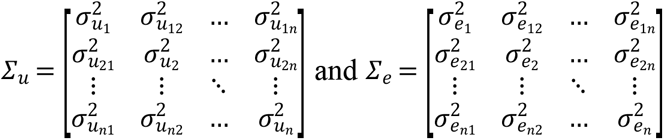

where 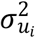 and 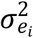 are, respectively, the additive and residual genetic variance associated with the *i^th^* trait with *i* = 1,…, *n* = 5; 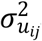 and 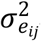 are, respectively, the additive and residual genetic covariance associated with the *i^th^* and *j^th^* traits with *j* = 1,…, *n* = 5 and *i* ≠ *j*. The variance and covariance components were obtained via the restricted maximum likelihood method (REML). The additive genomic relationship matrix (G) was obtained as described by Van Raden [27] by the centralization of the matrix of markers:

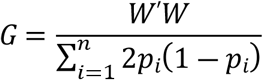

where, *W* is the centered marker matrix, which specifies the marker genotypes for each individual as 0, 1 or 2; *p_i_* is the frequency of the second allele at the *locus*.

With the genetic values of the individuals for the NB and MFB traits, the fruit production per plant (FPP) for each genotype was estimated.

The heritability in the strict sense of the *i^th^* trait was estimated following the equation below:

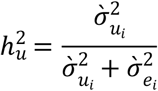

where 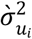 and 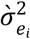 are, respectively, the additive genetic variance and the residual variance of the *i^th^* trait. The genetic correlation between the *i^th^* and *j^th^* traits was obtained through the following equation:

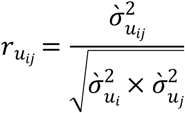

where 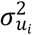 and 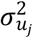 are, respectively, the additive genetic variance of the *i^th^* and *j^th^* trait, and 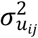 is the additive genetic covariance between the *i^th^* and *j^th^* traits.

### 2.7. Predictive ability and cross-validation

The predictive ability 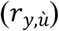 and the standard error of 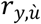, were estimated through the cross-validation procedure, subdividing randomly the population into 5 folds. Thus, 220 genotypes were used for training set and 55 genotypes were used for validation set. For each fold, the 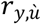 was obtained by the correlation between the predicted GEBV’s 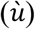 and the corrected phenotypic values (*y*).

### 2.8. Genetic selection based on selection index

With the GEBV’S predicted by the G-BLUP ST method, the selection of the best genotypes was carried out based on multiple variables. For this, the method of Mulamba and Mock (1978) was used, which is based on the sum of the individual ranks of each trait, creating a global rank. For the selection, the parameters of NB, MFB, PFP, RL and PY in the positive direction and EDF in the negative direction were considered. Two processes were performed before rank summing. The first was to transform the EDF values into classes, in order to create four classes. Class I for small fruits (values below the first quartile of the distribution of the genetic values); class II for small/medium size fruits (genotypes with values between the first and second quartile of the distribution of the genetic values); class III for medium fruits (genotypes with values between the second and third quartile of the genetic values), and class IV for large fruits (genotypes with values above the third quartile of the genetic values). The second change was the normalization of the number of ranks for each trait, to avoid traits with fewer ranks having a greater influence on the selection process. The standardization followed the expression below:

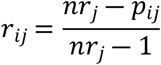

where *r_ij_* is the standardized rank value for genotype *i* and characteristic j; *nr_j_* is the number of ranks of trait *j* and *p_ij_* and is the rank of genotype *i* for the trait *j*.

In order to compare the selective efficiency between the G-BLUP ST and G-BLUP MT models, was estimated the expected selection gain for the population of the next selection cycle and for the commercial seed donor population. For both methods and all characteristics, with the exception of PFP, estimates were obtained by:

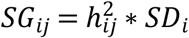

where, *SG* is the expected selection gain for characteristic *i* by model *j*; 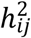 is the heritability of characteristic *i* estimated by model *j* and *SD_i_* is the selection differential for characteristic i in model j estimated based on corrected phenotypic values.

The SG for PFP was estimated based on the product of the SG estimates of NB and MFB.

### 2.9. Comparison between methodologies

Cohen’s Kappa coefficient (Cohen, 1960) was used to analyze the agreement between the best selected individuals between the G-BLUP ST e G-BLUP MT models, for the commercial seed donor population (formed by the six best ranked individuals), and the second selective cycle population (formed by the 50 best ranked individuals). Cohen’s Kappa coefficient is given by:

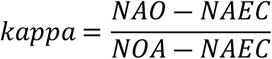

where *NAO* are the number of observed agreements, *NAEC* is the number of expected agreements by chance, and *NOA* is the number of analyzed observations (Resende et al., 2014).

## 3. Results

### 3.1. Data description

A summary of the descriptive statistics including mean, standard deviation, and maximum and minimum values for the five agronomic traits evaluated in this work is shown in Table 1. It is noted that in general, the population presented a small variation in the average phenotypic response between the years evaluated.

**Table 1.** Amplitudes, mean, standard deviation (SD) and coefficient of variation (CV) for rachis length (RL), fruit mass per bunch (MFB), number of bunches (NB), equatorial fruit diameter (EDF) and pulp yield (PY) for the years 2018, 2019, 2020 and 2021.

| Var. | Year | Minimum | Maximum | Average | SD | CV (%) |
| --- | --- | --- | --- | --- | --- | --- |
| **RL (cm)** | 2018 | 32 | 99 | 60.6 | 14.35 | 23.68 |
|  | 2019 | 33 | 104 | 62.72 | 14.15 | 22.56 |
|  | 2021 | 34.3 | 90.3 | 59.31 | 10.39 | 17.52 |
| **MFB (kg)** | 2018 | 0.4 | 8.3 | 3.05 | 1.57 | 51.47 |
|  | 2019 | 0.5 | 8.7 | 3.45 | 1.49 | 43.19 |
|  | 2021 | 0.3 | 8.7 | 3.68 | 1.87 | 50.81 |
| **NB** | 2018 | 1 | 7 | 4.24 | 1.35 | 31.84 |
|  | 2019 | 0 | 8 | 3.28 | 1.36 | 41.46 |
|  | 2020 | 0 | 7 | 3.69 | 1.03 | 27.91 |
|  | 2021 | 0 | 7 | 2.18 | 1.36 | 62.38 |
| **EDF (mm)** | 2018 | 11.15 | 18.9 | 14.03 | 1.15 | 8.20 |
|  | 2019 | 10.36 | 21.3 | 13.24 | 1.09 | 8.23 |
|  | 2021 | 10 | 19.94 | 13.31 | 1.25 | 9.39 |
| **PY (%)** | 2018 | 4.57 | 50.81 | 30.62 | 6.5 | 21.23 |
|  | 2019 | 16.75 | 51.52 | 35.96 | 5.37 | 14.93 |
|  | 2021 | 12.55 | 46.63 | 29.76 | 6.06 | 20.36 |

### 3.2. Genetic parameters, Genetic correlation and phenotypic correlation

The heritability and the environmental and additive genetics variance components were estimated for G-BLUP ST and G-BLUP MT models (Table 2). For both approaches, the higher residual variance compared to the variance due to genetic effects is emphasized, with the exception of EDF in the G-BLUP ST, where these estimates were equal (Table 2). For the traits evaluated, with RL exception, the G-BLUP ST model presented estimates of the genetic variance components 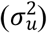 higher than the G-BLUP MT (Table 2).

**Table 2.** Additive genetic variance 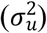, residual variance 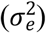 and restricted heritability 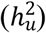 considering single trait (ST) and multiple trait (MT) models in genomic data.

| Model | Par. | RL | MFB | NB | EDF | PY |
| --- | --- | --- | --- | --- | --- | --- |
| **Single-trait** | $\sigma_u^2$ | 59.55 | 1.04 | 0.32 | 0.81 | 19.71 |
| | $\sigma_e^2$ | 73.53 | 1.15 | 0.72 | 0.41 | 4.96 |
| | $h_u^2$ | 0.45 | 0.48 | 0.31 | 0.67 | 0.80 |
| **Multi-trait** | $\sigma_u^2$ | 61.75 | 0.90 | 0.31 | 0.78 | 19.49 |
| | $\sigma_e^2$ | 72.17 | 1.21 | 0.73 | 0.41 | 5.06 |
| | $h_u^2$ | 0.46 | 0.43 | 0.29 | 0.65 | 0.79 |

Parameters (Par.); Rachis Length (RL); fruit mass per bunch (MFB); number of bunches (NB); equatorial diameter of fruit (EDF) and pulp yield (PY).

Comparing both models in relation to 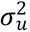, the G-BLUP ST provided estimates 15.56% (MFB), 3.23% (NB), 3.85% (EDF) e 1.13% (PY) higher than the estimated by G-BLUP MT. Due to these slight differences, and the similarity of the 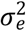 observed between the G-BLUP MT and G-BLUP ST, in general, the 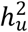 were similar between the tested models, ranging from 0.01 to 0.05 magnitude. The most significant difference was observed for MFB, which obtained a 0.05 higher 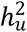 for the G-BLUP ST (Table 2).

The genetic and phenotypic correlations between the traits are shown in Fig. 2, which were calculated using the (co)variance estimates obtained by the G-BLUP MT model, so that the genetic correlation estimates are presented on the upper diagonal and, on the lower diagonal, estimates of phenotypic correlations are presented.

**Fig. 2.**
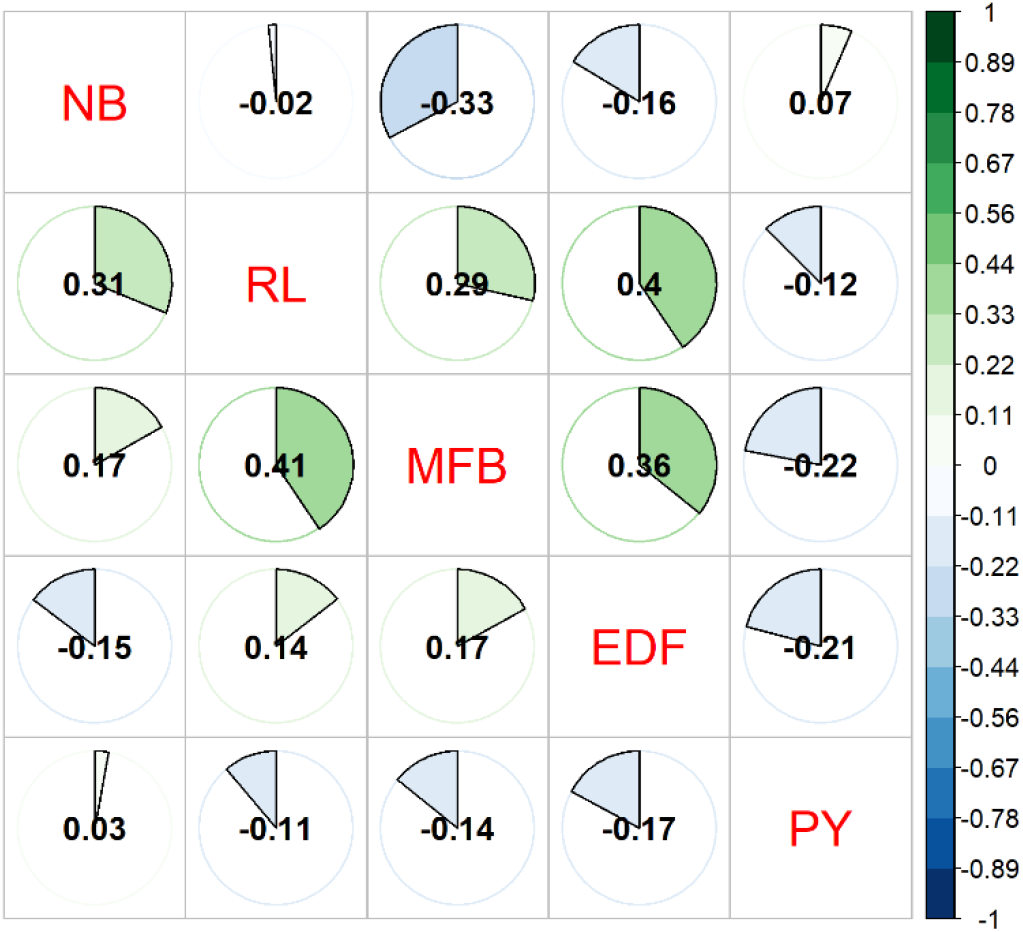
Genetic (upper diagonal) and phenotypic (lower diagonal) correlation between the traits RL (Rachis Length), EDF (Equatorial Fruit Diameter), MFB (Fruit Mass per Bunch), NB (Bunch Number) and PY (Pulp yield).

Comparing the phenotypic and genotypic correlations between the traits in Fig. 2, it is observed that only two correlation estimates had changes in their directions (RL and NB; MFB and NB), with the change in correlation between MFB and NB being more pronounced. The results obtained for the phenotypic and genotypic correlations can be classified from low to moderate (Evans et al., 1996). For phenotypic correlation, the highest values observed were 0.41 (RL and MFB), in the positive direction, and −0.17 (EDF and PY), in the negative direction. While, for the genotypic correlations between traits, the highest values were 0.40 (RL and EDF) in the positive direction, and −0.37 (NB and MFB) in the negative direction (Fig. 2).

### 3.3. Predictive ability

The predictive ability 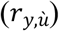 for the G-BLUP ST and G-BLUP MT models is shown in Table 3. For the G-BLUP ST, the 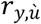 ranged from 0.21 for NB to 0.49 for PY, and for the G-BLUP MT model, the 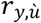 ranged from 0.18 for NB to 0.48 for PY. In general, the values of 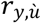 observed for both methodologies were close, as well as NB and PY maintained their positions with the lowest and highest 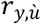, respectively. The results show that both models are similar to perform the prediction of genomic genetic values.

**Table 3.** Predictive ability 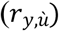 of the single-trait (G-BLUP ST), multi-trait (G-BLUP MT) models for RL (Rachis Length), EDF (Equatorial Fruit Diameter), MFB (Mass of Fruit per Bunch), NB (Number of Bunch) and PY (Pulp Yield).

| Parameters | G-BLUP ST | G-BLUP MT |
| --- | --- | --- |
| RL | 0.29 ($\pm 0.03$) | 0.22 ($\pm 0.01$) |
| MFB | 0.27 ($\pm 0.03$) | 0.26 ($\pm 0.04$) |
| NB | 0.21 ($\pm 0.06$) | 0.18 ($\pm 0.05$) |
| EDF | 0.44 ($\pm 0.02$) | 0.42 ($\pm 0.01$) |
| PY | 0.49 ($\pm 0.06$) | 0.48 ($\pm 0.07$) |

### 3.4. Matrix selection and expected genetic advancement

The Cohen’s Kappa coefficient was calculated for the top 20% of individuals (50) with the highest rank positions based on the Mulamba and Mock index (Mulamba and Mock, 1978), aimed to form the population of the second selection cycle. The agreement between the G-BLUP ST and G-BLUP MT models was 0.68 and is classified as substantial (0.60-0.80) (Sergio R. Munoz and Bangdiwala, 1997) (Fig. 3). However, when we reduce the selected population to six individuals, to build the donor population of commercial seeds, the coefficient is reduced to an agreement of 0.50, considered moderate.

**Fig. 3.**
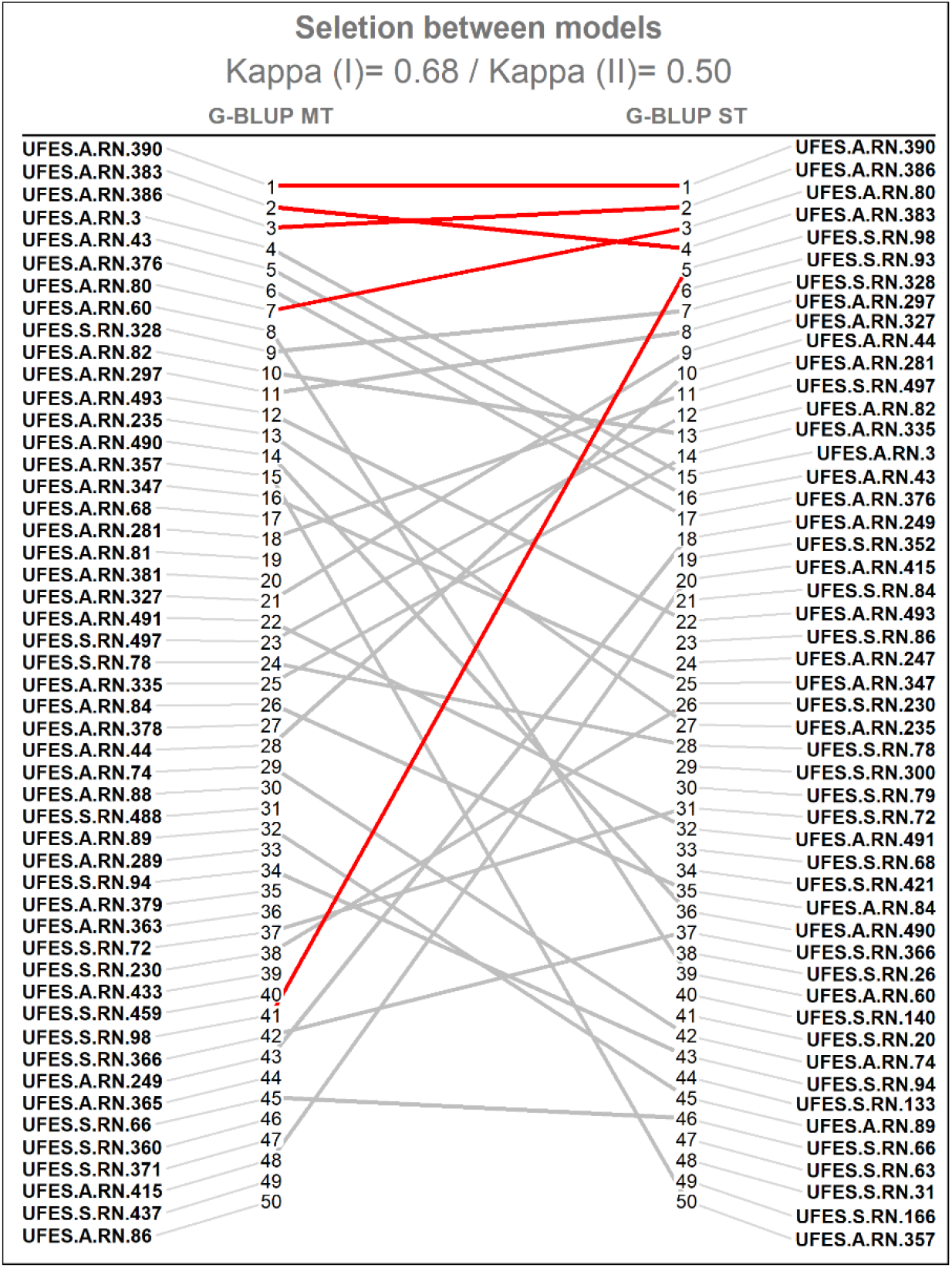
The Cohen’s Kappa coefficient showing the agreement between genotypes selected by the G-BLUP MT and G-BLUP ST models. Kappa (I) − 20% of the best genotypes (50 genotypes) and Kappa (II) - the six best genotypes selected.

Even though the agreements of the Cohen’s Kappa coefficient were classified from moderate (0.41-60) to substantial (0.61-60) (Munoz and Bangdiwala, 1997), is observed in Fig. 3, a great divergence in the ranking of individuals between the methods used. To support the comparison of the efficiency between the methods, the selection gain (SG) estimate was obtained through the heritability and phenotypic means of the selected individuals, and the results are shown in Table 4. In general, higher SG resulting from the G-BLUP ST model is observed for both populations.

**Table 4.** Estimates of selection gains provided by the single-trait (G-BLUP ST) and multi-trait (G-BLUP MT) models for the selected genotypes (S.I.) for the population of the next selection cycle (50 genotypes), and the commercial seed donor population (six genotypes).

| Trait | Population of the next selection cycle |  | Deed donor population |  |
| --- | --- | --- | --- | --- |
|  | G-BLUP ST | G-BLUP MT | G-BLUP ST | G-BLUP MT |
| **RL** | 31.25 | 31.64 | 30.58 | 32.58 |
| **MFB** | 2.06 | 1.62 | 2.40 | 1.76 |
| **NB** | 1.16 | 1.14 | 1.25 | 1.22 |
| **EDF** | 8.75 | 8.57 | 8.62 | 8.32 |
| **PY** | 27.60 | 27.10 | 30.15 | 29.86 |
| **PFP** | 2.39 | 1.85 | 3.00 | 2.15 |

RL: Rachis Length, EDF: Equatorial Fruit Diameter, MFB: Mass of Fruit per Bunch, NB: Number of Bunch, and PY: Yield of Pulp.

The population mean, the phenotypic and genotypic behavior of the selected individuals, based on the information from the G-BLUP ST model and the field phenotypic, is shown in Fig. 4. It is possible to observe that the phenotypic means of the six selected individuals, in the desired direction, have great performance in relation to the population average, being the improvement 31.13%, 51.05%, 98,40%, 11.56%, 15.57%, and 7.43% for NB, MFP, PFP, RL, PY and EDF, respectively.

**Fig. 4.**
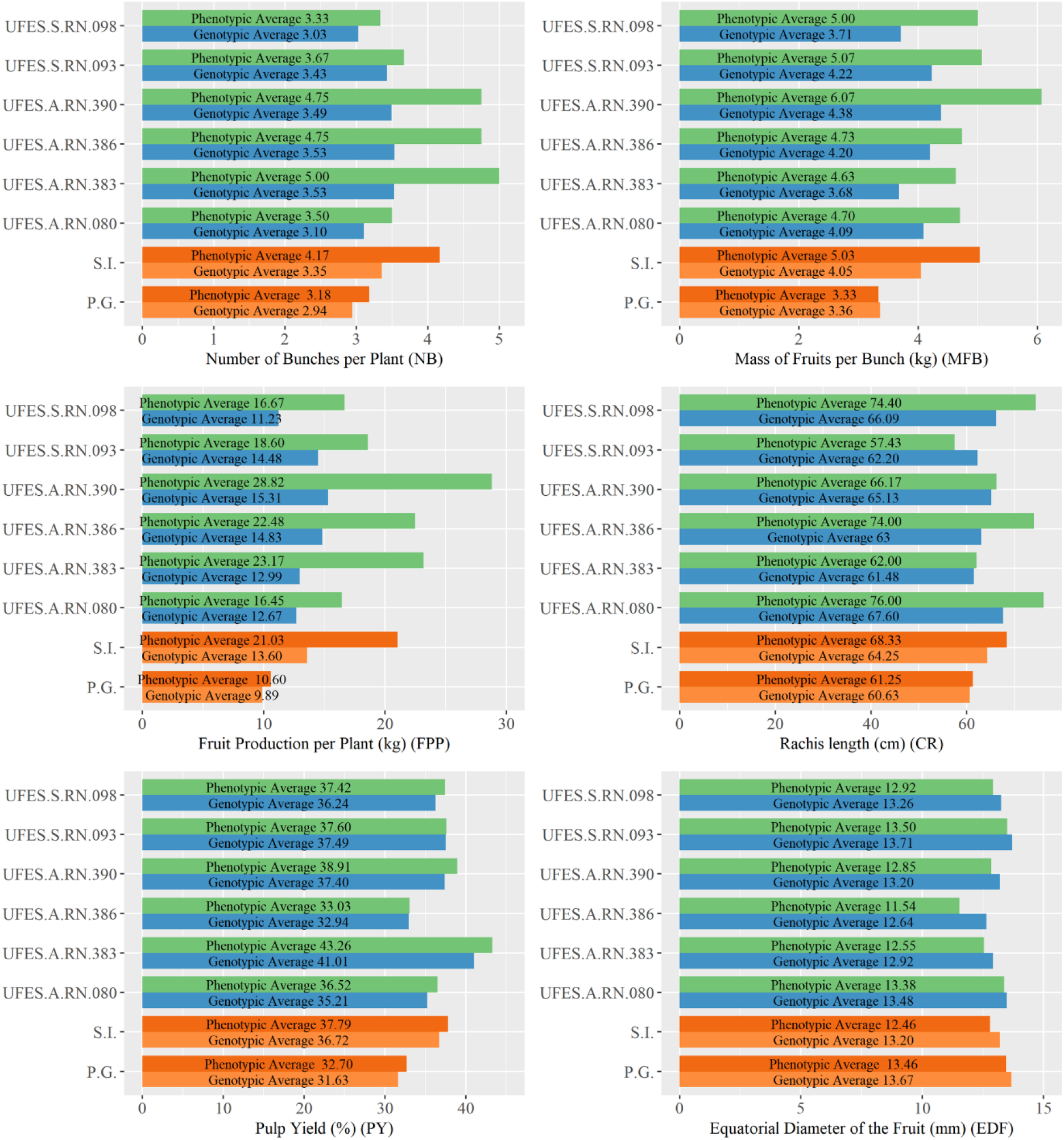
Phenotypic means observed in the field and GEBV’s + mean of selected genotypes and general population. S.I.: commercial seed donor population, P.G.: general population.

The selection of the six best genotypes provides a change in phenotypic responses equivalent to 0.99 bunches, 1.70 kg, 10.43 kg, 7.08 cm, 5.09%, and −1.00 mm for the traits NB, MFP, PFP, RL, PY and EDF, respectively (Fig. 4). With SG, is expected an improve on the averages in 0.41 bunches, −0.69 kg, 3.62 cm, 5.09%, and −0.47 mm, for NB, MFP, RL, PY, and EDF, respectively. For PFP, the genomic values were estimated by the product of the genomic values of MFB and NB, and the difference between the means of the selected and the general population was 3.71 kg (Fig. 4).

## 4. Discussion

As there are few programs that are based on the improvement of juçaizeiro, also a few scientific works that are aimed at the application of selective techniques for individuals of this species. In this sense, the existing breeding programs are in the early stages of development, and thus, their techniques are basically based on classical methodologies with phenotypic information. In contrast to these processes, the present work innovates in being the first scientific study focused on genomic selection of *Euterpe edulis*, with the objective of increasing selective accuracy and selection gains within the breeding cycle.

### 4.1. Genetic parameters, genetic correlation and phenotypic correlation

The 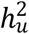 found in the present work for the traits under study indicate that the morphometric traits of fruits (EDF and PY) have a greater potential for heredity and genetic control, when compared to the other productive parameters related to clusters (CR, MFB and NB). This behavior was also observed in *Euterpe oleracea* (açaizeiro), a species of the same genus as juçaizeiro (Farias Neto et al., 2008; Teixeira et al., 2012).

It is expected that traits with higher 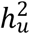, consequently, have a higher predictive ability (Legarra et al., 2008). The same was found by Legarra et al. (2008), where the traits with the highest 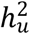 in the present study (EDF and PY) also had the higher predictive ability (Table 3). The higher values of 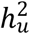 for the traits of EDF and PY in relation to NB, MFB and RL can demonstrate the dynamism of the behavior of the 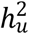 as a function of the hereditary and environmental behavior. It is known that for quantitative traits, several environmental factors can affect phenotypic behavior. In this sense, it can be assumed that the genetic control of NB, MFB and RL is reduced due to the exposure of these traits for a longer period of time to environmental effects and more variable conditions that are not controlled, for example, direction of insertion of the bunch into the matrix plant, exposure to inclement weather, feeding the fauna, among others. While, the development of fruits in the bunch will occur, for example, in a fraction of the time of the infructescence development cycle, which will be located in the same region of the plant, with a similar influence on luminosity, temperature, and availability of nutrients. Therefore, genetic control can be expected to be greater.

Published scientific studies evaluating associations between *Euterpe edulis* traits are scarce in the literature (Marçal et al., 2015; Oliveira et al., 2015; Silva et al., 2018). However, these analyzes are fundamental for understanding the interactive behavior between traits, revealing linear cause and effect responses that are essential for the agricultural development of the crop and for breeding programs, as it allows the determination of different types of practical strategies.

The knowledge of correlations in the genetic scope is important in breeding programs because they are heritable and can be used in indirect selection (Cruz et al., 2012). However, the evaluation of phenotypic behavior is also essential, given the fact that traits genotypically and non-phenotypically associated end up not having practical applicability in the indirect selection process, since the phenotype is influenced both by genetic and environmental effects.

The (co)variances estimated by the G-BLUP MT, made it possible to estimate the Pearson correlation for genetic and phenotypic effects between the traits RL, EDF, MFB, NB and PY (Fig. 2), allowing a counterpoint between these estimates. In this sense, we noticed that practically all the associations preserved their influence on the behavioral response, with the exception of the NB-RL and NB-MFB pairs, which had an inversion of the sense of their estimates. In this condition, indirect selection is compromised by the fact that selection by genotypic responses may not be expressed in the desired sense in the environment. With this, we can conclude that the analyzes of phenotypic and genotypic associations are complementary and make it possible to determine practical actions, using indirect processes to enable increased gains in breeding programs.

Evaluating biometric characteristics of juçaizeiro fruits, Marçal et al. (2015) reported a positive genetic association between EDF and fresh seed mass (0.75). Knowing that the juçaizeiro fruit is mostly composed of seed, and the pulp makes up a small portion of it, increasing EDF can reduce the percentage of pulp yield. This fact is confirmed by the association observed in Fig. 2, which shows inversely proportional associations between EDF and PY (−0.17 phenotypic; −0.21 genetic), that is, the increase in fruit size results in a reduction in the percentage of pulp per fruit, harmful to industrial processing.

The results observed by Oliveira et al. (2015)for the genetic correlation in *Euterpe edulis*, are close to those observed in the present work between the characteristics NB, PY and RL. In both situations, the magnitudes of the associations can be classified from very weak to weak (Evans et al., 1996). Oliveira et al. (2015) reported associations of 0.20, −0.10 and −0.02 for NB-PY, PY-RL and NB-RL, respectively, while the present work obtained estimates of 0.07, −0.12 and −0.02, respectively. Corroborating these results, Farias Neto et al. (2016) also found a weak correlation between NB-RL (−0.01) for *Euterpe oleracea*.

Regarding the correlation results observed in Fig. 2, we can determine that the use of indirect selection among the evaluated characteristics would be inefficient. Therefore, the effect that the characteristics presented among themselves were mostly low, making the use of this practice unfeasible due to the small gains that could be obtained.

### 4.2. Predictive ability

In order to increase selection efficiency, choosing the best methodology for analyzing and predicting the genetic values of individuals is one of the fundamental aspects to be taken into account in a breeding program. In this study, we compare G-BLUP ST e G-BLUP MT models for the evaluation of genomic prediction in a base population of juçaizeiro. The existence of genetic correlations between the selection traits is the basis for the advantages presented by G-BLUP MT (Jia and Jannink, 2012) and the absence of correlation could lead to equivalent or even superior results when using G-BLUP ST (Jia and Jannink, 2012). Therefore, we can conclude that, depending on the analyzed traits and the existence of correlations, it is expected that the use of the GBLUP MT model will increase the accuracy of the evaluation predictions (Tsuruta et al., 2011).

However, the superiority of G-BLUP MT over G-BLUP ST for 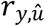 was not observed in the present work. The results obtained by the G-BLUP MT model showed similar values of 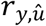 when compared to the G-BLUP ST model, which may have been caused by the linear associations that varied from low to moderate intensity (≤ 0.42) ( Fig. 2). Therefore, the present scenario of low association between the characteristics evaluated initially suggests the advantage that the use of genomic selection can provide to multi-trait models. As reported by Calus and Veerkamp (Calus and Veerkamp, 2011), who observed an improvement in 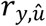 when marker information was added to the model for the traits that had associations less than 0.50. In this sense, it is expected that the flow of genetic and phenotypic information provides higher 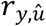 when compared to G-BLUP ST.

The slightly lower performance of G-BLUP MT for predictive abilities can be explained by the low genetic and residual correlations between traits, as reported by Runcie and Cheng (2019), who attribute such results to the fact that low correlations can cause imprecision in the estimates of genetic and residual covariance parameters may result in reduced model performance. However, as the 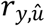 were close, there is no evidence of such negative effects on the covariance estimates, deducing that the flow of information between the characteristics was not sufficient to improve the 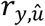 of the model G-BLUP MT in relation to the G-BLUP ST, considering that marker information was used by both models.

For the five characteristics evaluated, the biggest difference between the 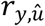 for the methods tested was for RL, which presented values of 0.29 and 0.22, for the G-BLUP-ST and G-BLUP MT, respectively. The superiority of the G-BLUP-ST model diverges from patterns observed by other studies (Arojju et al., 2020; Bhatta et al., 2020; Guo et al., 2014; Jia and Jannink, 2012; Okeke et al., 2017; Rutkoski et al., 2012; Tsai et al., 2020; Tsuruta et al., 2011), in which superiority is observed for the G-BLUP MT model. As it was designed to benefit from the existence of genetic correlations between traits, it is expected that the G-BLUP MT model will present better results than the G-BLUP ST (Calus and Veerkamp, 2011; Schulthess et al., 2016). Due to the similarity of the results of the predictive ability of the models G-BLUP ST and G-BLUP MT, it is not possible, just for this parameter, to indicate the best model for the selection of juçaizeiro genotypes.

### 4.3. Matrix selection and expected genetic advancement

To build the second selective cycle breeding population, in order to maintain genetic variability, a selection intensity of approximately 20% was defined. In this condition, the Kappa agreement observed among the approaches evaluated was classified as substantial (0.68), considering that its value was between 0.60 and 0.80 (Munoz and Bangdiwala, 1997). This similarity between the approaches can be explained by the fact that as the variables under study present low to moderate correlations (Evans et al., 1996), both methods ended up leading to similar results.

We should add that the high Kappa agreement between the methods is also associated to the conditions of the phenotypic data set used in the study, which has a low degree of imbalance. Thus, the predictive abilities of the approaches did not have a great impact on the selection of different genotypes between models; however, the influence on the ranking of the best individuals was strongly impacted, as shown in Fig. 3.

When considering the six best genotypes, the selection of the best genotypes had greater divergence between the models, in which the Kappa value was 0.50, evidencing the need to determine the best model to carry out the selection process, aiming at greater future gains. In this way, the models are chosen according to their ability to predict more accurately the GEBV’s of the analyzed individuals. However, in the present study, 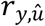 was not sufficient to determine the best method, due to the similar estimates observed.

Due to the similarities of the 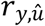 and the great divergence in the ranking for the Kappa agreement, the choice of the best model for the selection of the juçaizeiro genotypes was based on the SG. In order to standardize the comparison between the models and eliminate the differences in the GEBVs associated with the form of prediction, the SG of each trait was estimated by the corrected phenotypic means and heritability from each model for the populations selected for the next selection cycle and seed donors. In this sense, the G-BLUP ST was chosen for the selection of juçaizeiro genotypes, because this model provided higher SG, in addition to being a simpler and more usual method for breeding programs.

The production of fruits per plant is a feature that serves both the productive sector and the industrial sector, as it provides greater economic return to producers and increases the amount of raw material for industries that currently suffer from a shortage of this base product for their operations. In this sense, observing the difference in selection by phenotypic values, the change in the average fruit production per plant has an increase of 9.67 kg, which indicates a great advance for the culture due to the possibility of achieving great gains in productivity, resulting in an incentive to install of culture.

Due to the juçaizeiro is a wild and perennial species, which presents a series of characteristics intrinsic to its development, the practical activities of conventional breeding techniques become more difficult. In general, when evaluating the structure of a breeding program aimed at juçaizeiro, it is expected that the gains obtained per unit of time will be reduced. In this sense, the present study stands out for applying genomic selection techniques in *Euterpe edulis*, a wild open-pollinated species, aiming to increase gains per unit of time.

Therefore, perhaps the biggest advantage brought by this methodology for a wild breeding population, as is the case of *Euterpe edulis*, is the use of the genomic matrix applied to the relationship correction. Therefore, the insertion of this information provides more accurate estimates and, consequently, greater reliability to the program. Furthermore, the use of genomic selection may allow a reduction in the selection cycle by being able to predict the behavior of non-evaluated genotypes.

All the traits evaluated in the study influence the economic potential of the species, covering the interests of the productive sector and the industrial sector. Additionally, in function of the alterations observed by the selection of the six superior genotypes, we can conclude that the genomic prediction using the G-BLUP ST was efficient to provide alterations of the means in the desired directions. That is, obtaining an increase in the average phenotypic response of the selected genotypes for the characters of NB, MFB, PFP and PY, and a reduction for EDF.

## 5. Conclusion

Our results showed that the GBLUP-ST genomic prediction was more efficient in selecting the best genotypes, so that the selection provided substantial gains in the desired direction for the multiple traits. Thus, the six selected genotypes (UFES.A.RN.390, UFES.A.RN.386, UFES.A.RN.080, UFES.A.RN.383, UFES.S.RN.098 e UFES.S.RN.093) can be used as commercial seed donor matrices for the development of seedlings for the implantation of productive orchards, which will meet the demands of the productive sector and the consumer market.

## Declaration of Competing Interest

The authors declare that they have no known competing financial interests or personal relationships that could have appeared to influence the work reported in this paper.

## Acknowledgments

We thank the Conselho Nacional de Pesquisa (CNPq, Brazil) (Researcher productivity fellowship AF; MFSF #311840/2019-1), Coordenação de Aperfeiçoamento de Pessoal de Nível Superior (CAPES, Brazil) – Finance Code 001, and the Fundação de Amparo à Pesquisa do Espírito Santo (FAPES, Vitória – ES, Brazil) in partnership with VALE, for the financial support to this study. We thank the companies, Açai Juçara and Bonaloti, for supporting research development and we are also grateful to Pedro and Vicente Bortoloti, the owners of the managed area. We would like to thank all colleagues at the Biometrics and Genetics and Plant Improvement laboratories from the Universidade Federal do Espírito Santo, for helping to process and organize the samples.

## CRediT authorship contribution statement

**Guilherme Bravim Canal:** Conceptualization, Methodology, Formal analysis, Investigation, Data Curation, Writing - Original Draft, Writing - Review & Editing. **Cynthia Aparecida Valiati Barreto:** Methodology, Formal analysis, Data Curation, Writing - Original Draft, Writing - Review & Editing. **Marcia Flores da Silva Ferreira:** Conceptualization, Resources, Writing - Review & Editing, Funding acquisition. **Camila Ferreira Azevedo:** Conceptualization, Methodology, Writing - Review & Editing. **Moysés Nascimento:** Conceptualization, Methodology, Writing - Review & Editing. **Francine Alves Nogueira de Almeida:** Data Curation, Methodology. **Diego Pereira do Couto:** Data Curation, Methodology. **Iasmine Ramos Zaidan:** Data Curation, Methodology. **Adésio Ferreira:** Conceptualization, Methodology, Resources, Formal analysis, Investigation, Writing - Review & Editing, Funding acquisition, Project administration.

## Data availability statement

The datasets generated during and/or analysed during the current study are available from the corresponding author on reasonable request.

## Notes

### Competing Interest Statement

The authors have declared no competing interest.

